# Accelerated neural aging as a systems-level vulnerability for tinnitus

**DOI:** 10.64898/2026.05.13.724786

**Authors:** Fabian Schmidt, Gianpaolo Demarchi, Nadia Müller-Voggel, Seyed Mostafa Kia, Eugen Trinka, Nathan Weisz

**Affiliations:** Paris-Lodron-University of Salzburg, Department of Psychology, Centre for Cognitive Neuroscience, Salzburg, Austria; Department of Neurosurgery, Universitätsklinikum Erlangen, Friedrich-Alexander University Erlangen-Nürnberg; Center of Cognitive Science and Artificial Intelligence, Tilburg University, Tilburg, the Netherlands; Donders Institute for Cognition, Brain and Behavior, Radboud University, Nijmegen, The Netherlands; Department of Psychiatry, UMC Utrecht Brain Center, University Medical Center, Utrecht, the Netherlands; Neuroscience Institute, Christian Doppler University Hospital, Centre of Cognitive Neuroscience, Paracelsus Medical University, Salzburg; Department of Neurology, Neurocritical Care and Neurorehabilitation, Christian Doppler University Hospital, Centre of Cognitive Neuroscience, Paracelsus Medical University, Salzburg, Member of EpiCARE, Austria

## Abstract

Tinnitus, the perception of sound without an external source, affects 10-15% of individuals, yet its neural mechanisms remain poorly understood. Building on evidence that chronological age predicts tinnitus risk beyond hearing loss, we tested the hypothesis that accelerated neural aging increases susceptibility to tinnitus by integrating cross-sectional resting-state MEG with longitudinal structural imaging data. In a large MEG dataset, spectral parametrization revealed that age-neural relationships were amplified in the tinnitus group compared to age-, sex- & hearing-matched controls. Complementing these findings, prospective analyses using data from the UK Biobank showed that among participants without tinnitus at baseline, stronger age-related declines in white-matter density predicted later tinnitus onset. Collectively, these converging functional and structural findings support accelerated brain aging as a key risk factor for tinnitus.

## Introduction

Approximately 10–15 % of individuals report the perception of sound in the absence of any external source, a phenomenon known as tinnitus [1]. Typically, tinnitus presents itself as simple auditory sensations (most often pure tones or narrow-band noise) and is thus phenomenologically distinct from the complex auditory hallucinations observed in psychiatric disorders. Clinically significant distress arises in roughly 1–3 % of all people, yet no interventions reliably target its underlying causes. Despite affecting a substantial proportion of the population, the etiological factors and neural mechanisms driving both the onset and persistence of tinnitus remain poorly understood. A central challenge in tinnitus research is that the conscious auditory experience cannot yet be objectively quantified using measures of brain structure or function. As a result, both assessment and the development of targeted interventions still rely exclusively on subjective self-report. Furthermore, despite individuals sharing similar risk factors, current research is unable to predict who will develop tinnitus. Here, we confront both challenges by integrating a cross-sectional analysis of spontaneous brain activity with a longitudinal assessment of structural brain changes. Specifically, we test the hypothesis that accelerated neural aging constitutes a risk factor for tinnitus [2, 3] by investigating whether age-related neural signatures are amplified in affected individuals.

Many researchers and clinicians in the field of tinnitus propose that tinnitus arises from aberrant brain activity triggered by cochlear injury [4]. By definition, tinnitus is the perception of sound in the *absence* of external stimulation, so most investigations have focused on spontaneous neural activity. Animal models of tinnitus consistently reveal abnormal—typically increased—spontaneous firing rates and enhanced synchrony in auditory processing regions, spanning from the brainstem [5, 6] through thalamic nuclei [7] to auditory cortex [8]. A leading hypothesis holds that such hyperactivity reflects increased gain or stochastic resonance in central auditory pathways following peripheral damage [9, 10]. In human studies, noninvasive electrophysiological techniques like EEG and MEG measure population-level signals rather than single-unit activity. Consequently, research has commonly compared band-limited oscillatory power between tinnitus and control groups, reporting alterations in delta, theta, alpha, and gamma bands [11, 12]. This focus is both pragmatic—given methodological constraints—and theoretically grounded in models linking aberrant oscillations to tinnitus: for example, enhanced delta/gamma activity in thalamocortical dysrhythmia [13], reduced alpha power reflecting cortical disinhibition [14, 15] as well as increased perceptual bias [16], and elevated gamma associated with perceptual binding and awareness [17]. Over two decades, a considerable number of EEG/MEG studies have identified intriguing patterns suggestive of altered excitatory/inhibitory (E/I) balance in tinnitus. Yet, a disappointing lack of reproducibility persists across studies [12], impeding both mechanistic understanding and clinical translation such as development of objective biomarkers or therapies (e.g., neurofeedback, brain stimulation) that target presumed oscillatory abnormalities [18, 19].

Also, the notion that hearing loss invariably leads to tinnitus is incomplete: 4–65% of individuals with hearing impairment do not develop tinnitus [20, 21]. This gap suggests that, beyond hearing loss as an enabling factor, additional contributors must be considered (see e.g., [22]). We have recently turned our attention to aging, which is associated with both increased tinnitus prevalence and hearing loss [23]. In three large cohorts using distinct metrics of auditory function, we demonstrated that chronological age predicts tinnitus risk over and above hearing loss [2, 3]. Here, age likely serves as a proxy for underlying neurobiological processes. Several age-related neural changes map onto proposed mechanisms of tinnitus—for example, shifts in excitatory/inhibitory balance, downregulation of inhibitory neurotransmission [24], and heightened reliance on predictive coding in perception [25]. On this basis, we previously speculated [2] that accelerated or advanced brain aging may be a critical factor in the emergence of tinnitus.

We combined large-N MEG resting-state data from our own lab with longitudinal structural imaging from the UK Biobank to address our hypotheses. Going beyond prior work (for reviews, see [12, 26]), we show that, by carefully parameterizing spectral activity [27], not only periodic effects—particularly in the alpha range—but also aperiodic features differ between individuals with tinnitus and controls. This pattern suggests that both components capture complementary aspects of tinnitus-related brain dynamics. Importantly, with regards to our neural aging hypothesis, the relationship between age and spectral measures is more extreme in the tinnitus group. Extending this cross-sectional dataset, among individuals without tinnitus at baseline, greater age-related declines in white-matter volume—measured prospectively in UK Biobank follow-up scans—predicted the subsequent onset of tinnitus. All these effects are robust even after rigorously matching participants on age, sex, and audiometric thresholds. Together, these findings support our hypothesis that accelerated or aberrant brain aging increases susceptibility to tinnitus.

## Results

Here we analyzed two large cohorts of tinnitus and carefully matched control participants (see **Figure 1A** **&** **Figure 3B**). Cohort#1 (*N* = 106) consists of eyes-open resting-state MEG recordings. To move beyond traditional band-limited power analyses (conflating periodic and aperiodic contributions), we applied data-driven spectral parameterization to separate periodic oscillatory peaks from broadband aperiodic activity [27]. Specifically, we used an IRASA-based approach [28, 29], to decompose each spectrum into periodic peak and aperiodic broadband parameters (e.g. spectral offset & exponent; see also Methods – Spectral Analysis). We then used linear mixed-effect models to identify which spectral parameters distinguished tinnitus from controls and to test whether these features showed differential age dependence between groups. Feature selection and the definition of age-sensitive metrics were informed by an independent lifespan dataset (CamCAN; [30]), which also provided a reference for normative age-related trajectories. Individuals with tinnitus in this sample, reported overall very low distress (79% compensated; see **Supplementary Table 1**). To directly assess aging-related vulnerability, Cohort#2 comprised longitudinal structural MRI from the UK Biobank (*N* = 356), enabling us to track changes in global, age-sensitive brain measures (defined again in CamCAN) over 2.73 years (*SD* = 1.33) in individuals who were tinnitus-free at baseline and matched for age, sex & hearing status at baseline and follow-up.

**Figure 1:**
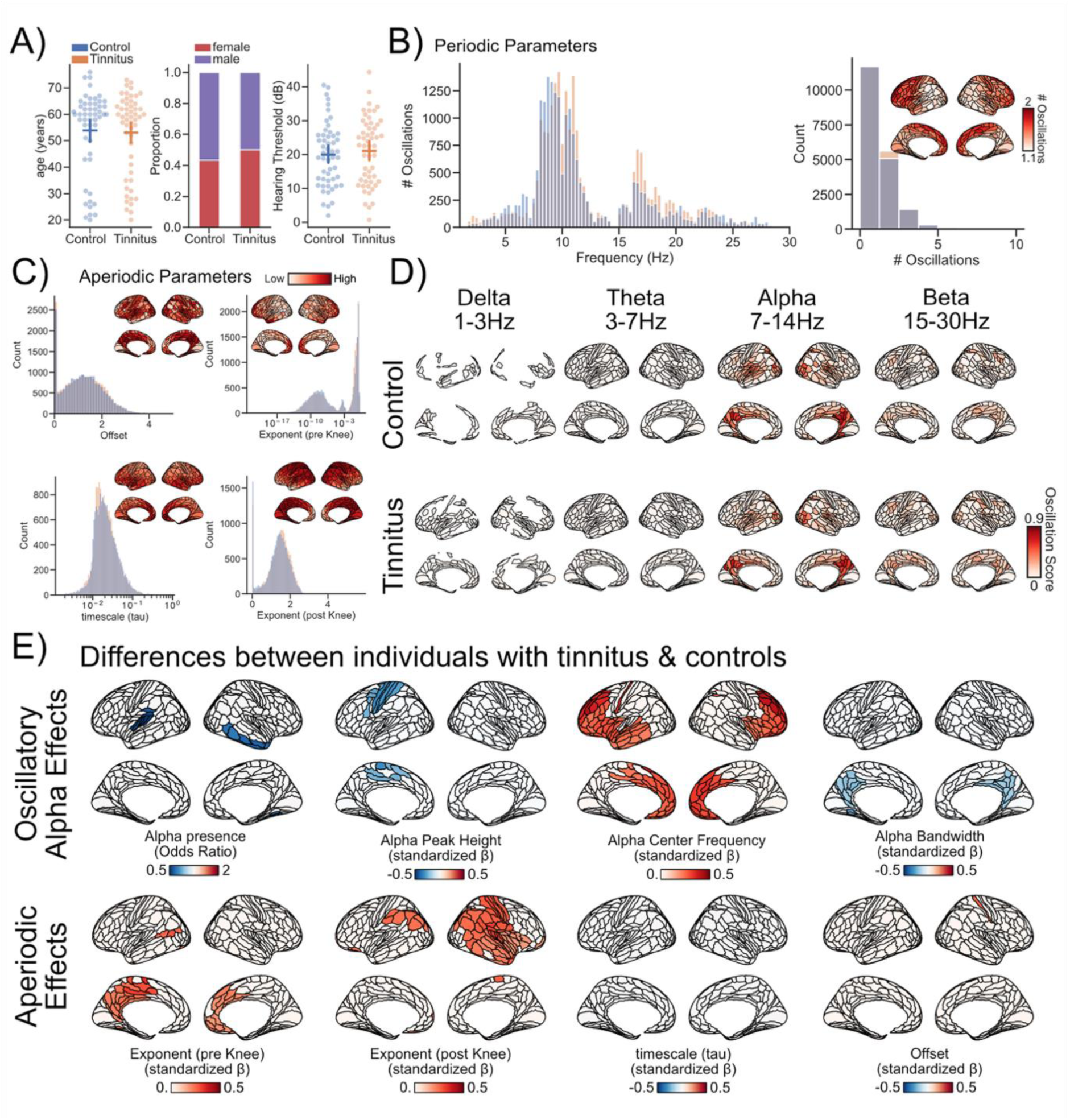
Periodic and aperiodic spectral features in tinnitus and controls. A) Resting state MEG data of *N* = 53 individuals with tinnitus (orange) and (*N* = 53) age, sex, and hearing matched controls (blue) were analyzed. Overall, patterns of periodic and aperiodic resting state activity are largely similarly distributed in individuals with tinnitus and matched controls (BCD). E) parcel-wise group differences (tinnitus vs control) for oscillatory alpha features (peak presence as odds ratio; peak height, center frequency, and bandwidth as standardized β coefficients and aperiodic features (offset, pre-knee exponent, post-knee exponent, timescale (τ). For all cortical maps in E, red colors indicate higher and blue colors lower values in tinnitus; color bars denote effect sizes for each parameter as mean standardized β coefficients. Coefficients where the 89% HDI overlaps with a region of practical equivalence are masked.

## Periodic and aperiodic spectral parameters associated with tinnitus

As tinnitus is a perception not elicited by external stimuli, most neuroscientists assume that perception-related neural activity patterns should be reflected in ongoing spontaneous activity. To counteract the low reliability of previous findings, we included a large and well-matched sample (**Figure 1A**; age: *β_standardized_* = -0.043, 89%HDI [-0.36, 0.28]; sex: *β_standardized_* _(log-odds)_ = -0.523, 89%HDI [-1.20, 0.17]; hearing threshold: *β_standardized_* = 0.098, 89%HDI [-0.22, 0.40]) and applied a spectral parameterization approach to source-level projected MEG data (see *Methods – Spectral Analysis*). This allowed us to disentangle periodic and aperiodic signal components. At a descriptive and more global level, the spectral and spatial distributions of aperiodic (**Figure 1C**) and periodic (**Figure 1BD**) components appear broadly similar between individuals with and without tinnitus (see also **Supplementary Figure S1**). Regarding the periodic parameters (**Figure 1B**), oscillatory peaks occur across a wide frequency range, with pronounced clusters in the classical alpha band (7–14 Hz) and the beta band (15–30 Hz), and smaller contributions in the delta (1–3 Hz) and theta (3–7 Hz) ranges (bands definition based on [27]). For these frequency bands, an oscillation score (quantifying the probability of a band-specific oscillation, weighted by relative band power, after adjusting for the aperiodic component; [27]) was computed indicating the consistency of oscillations first and foremost in the alpha band across tinnitus and control subjects (see **Figure 1D**). Overall, these patterns corroborate previous reports on oscillatory spectral features [27] and provide strong support for both the data quality of the present study and the validity of the spectral parameterization approach forming the basis for subsequent analyses.

While a descriptive inspection of the data suggests certain differences between individuals with and without tinnitus (e.g., a higher prevalence of peaks in the upper alpha range; **Figure 1B**), such visual impressions do not permit reliable inference about which resting-state spectral features are truly related to tinnitus. To address this, we employed linear mixed effect models to predict aperiodic and periodic parameters based on the presence or absence of tinnitus. Applying this modeling approach implicated multiple spectral features and spatially distinct brain regions (**Figure 1E**).

As a previously unreported finding, we observed a steeper decay of power with frequency in individuals with tinnitus both for the pre- and post-knee exponents (i.e., increased spectral exponents). Spatially, pre-knee exponent differences were mainly localized to medial areas, whereas post-knee differences were more widespread and predominantly lateral (**Figure 1E**) with a marked right-hemispheric lateralization. Across parcels, parcel-specific posterior mean effects for the pre-knee exponent averaged *β_standardized_* = 0.22 (range across parcels: 0.18–0.34), and for the post-knee exponent averaged *β_standardized_*= 0.23 (range: 0.17–0.31). In a previous study [31], investigating large-N samples, we observed that aperiodic age-related slope associations are also captured by cardiac activity. This suggests that cardiac activity could influence MEG aperiodic slope estimates through volume conduction. A control analysis, applying the same analysis procedure and statistical modeling to ECG data, indicated no comparable pre- or post-knee exponent differences between individuals with and without tinnitus (see **Supplementary Table 2 & 3**). This suggests that the reported aperiodic exponent effects are unlikely to be an epiphenomenon of differential cardiac activity.

Accounting for the influence of aperiodic components further revealed multiple periodic parameters in the alpha band that significantly predicted tinnitus status. We decided to focus on the alpha band due to its omnipresence both in tinnitus and controls and based on previous findings highlighting implications in tinnitus [14, 32]. Moreover, after accounting for aperiodic influences, only a small number of regions exhibited reliable oscillatory activity, this absence of oscillatory activity was most notably in the delta and theta range (**Figure 1D**; see also [33]). Notably, we observed a decrease in the probability that the spectral parametrization algorithm detected oscillatory activity in the alpha band, with the most pronounced effect in left auditory cortex (**Figure 1E**). Across parcels, posterior mean odds ratios averaged OR = 0.57 (range across parcels: 0.51–0.63), indicating ∼37–49% lower odds of alpha peak detection in the tinnitus group. Conditionally on alpha peak detection, we found that alpha peak amplitude was reduced in individuals with tinnitus, with effects that were predominantly left-lateralized and centered on sensorimotor and premotor regions (**Figure 1E**). Across parcels, parcel-specific posterior mean effects averaged *β_standardized_*= −0.26 (range across parcels: −0.33 to −0.22), indicating a small to medium reduction in alpha peak power in the tinnitus group. Furthermore, we observed a marked shift in the alpha range, such that tinnitus was associated with increased alpha peak (center) frequencies. Spatially, these effects were bilateral and predominantly frontal, with the strongest effects over frontal cortices and additional ventral/temporal involvement (see **Figure 1E**). Across parcels, parcel-specific posterior mean effects averaged *β_standardized_* = 0.28 (range across parcels: 0.21–0.42), consistent with a small to medium effect size.

When these peak frequency shifts and alpha power reductions are considered alongside the aperiodic contributions, the overall pattern suggests a dissociation between periodic and aperiodic dynamics in tinnitus. Reduced alpha peak detection and lower alpha peak height point to weakened alpha-band rhythmic activity, which is often interpreted as reduced oscillatory (alpha-mediated) inhibition and thus increased cortical excitability [34]. The accompanying increase in alpha center frequency is in line with computational work linking faster alpha rhythms to shifts in circuit excitability and inhibitory time constants [35]. By contrast, the steeper high-frequency aperiodic exponent has been proposed to reflect stronger inhibitory contributions in the broadband component [36, 37]. Taken together, these findings are compatible with decreased oscillatory inhibition in the alpha band alongside increased aperiodic inhibition, potentially reflecting distinct underlying processes.

Additionally, we observed a narrowing of alpha bandwidth that was largely parietal, suggesting reduced variability in rhythmic alpha activity in individuals with tinnitus (**Figure 1E**). Parcel-specific posterior mean effects averaged *β_standardized_* = −0.20 (range across parcels: −0.27 to −0.18), indicating a small reduction in alpha bandwidth in the tinnitus group. Neither neural timescales (τ) nor the spectral offset show notable group level differences between individuals with tinnitus and controls (aside from a small elevation of the spectral offset in somatosensory regions).

In sum, using a large sample and a spectral parameterization approach, we identify a rich constellation of alterations in spectral features—both periodic and aperiodic—in association with tinnitus, distributed across widespread brain regions. Importantly, these alterations are not just merely a consequence of the used parametrization approach, but are also visible in the raw unparametrized power spectra (see **Supplementary Figure S2**). Some of these effects align with established observations, including reduced alpha rhythmicity (lower alpha peak detection and peak height; [14]), whereas others extend prior work by revealing systematic shifts in alpha peak characteristics (increased center frequency and narrowed bandwidth, predominantly in fronto-parietal and parietal regions). Beyond periodic activity, tinnitus was also associated with changes in aperiodic parameters, especially steeper spectral exponents, suggesting that broadband and rhythmic processes may be differentially affected. Together, these results point to a broad reconfiguration of cortical dynamics in tinnitus, marked by diminished oscillatory alpha inhibition alongside distinct changes in the broadband aperiodic background activity.

## Spectral patterns of brain aging are more pronounced in individuals with tinnitus

Beyond examining group differences between individuals with and without tinnitus, the primary aim of our study is to test the hypothesis that aberrant neural aging trajectories may confer an increased risk of developing tinnitus. To this end, we first identified features of brain function that are strongly associated with chronological age, before assessing their relationship to tinnitus (using MEG data analyzed with the same source-level spectral parameterization approach). To ensure a clear separation between feature selection and group comparison, we used the *Cambridge Centre for Ageing and Neuroscience* (Cam-CAN) dataset as an independent reference. Specifically, we computed regression models between age and commonly associated spectral M/EEG parameters.

In line with previous MEG and EEG studies [38, 39], alpha peak (center) frequency decreased with increasing chronological age (**Figure 2A**). Across parcels, parcel-specific posterior mean effects averaged *β_standardized_* = −0.31 (range across parcels: −0.44 to −0.14), indicating a robust and widely distributed age-related slowing of alpha peak frequency. Among the aperiodic parameters, both exponents exhibited consistent associations with age: the pre-knee exponent decreased with increasing age (i.e., a flatter low-frequency slope), whereas the post-knee exponent increased (i.e., a steeper high-frequency slope). Across parcels, parcel-specific posterior mean effects for the pre-knee exponent averaged *β_standardized_* = −0.13 (range across parcels: −0.25 to −0.09), while effects for the post-knee exponent averaged *β_standardized_* = 0.21 (range across parcels: 0.1 to 0.38). The neural timescale (τ) also decreased with increasing age, indicating a shift towards shorter timescales in older participants. Across parcels, parcel-specific posterior mean effects averaged *β_standardized_* = −0.28 (range across parcels: −0.47 to −0.11). These periodic and aperiodic age effects were mostly bilateral and widespread with notable foci in temporal areas for the pre- and post-knee exponent and alpha center frequency changes. A schematic summary of these findings is presented in **Figure 2B**. Together, the results reveal widespread age-related changes in resting-state MEG activity, dominated by alterations in the aperiodic components of the spectrum.

**Figure 2.**
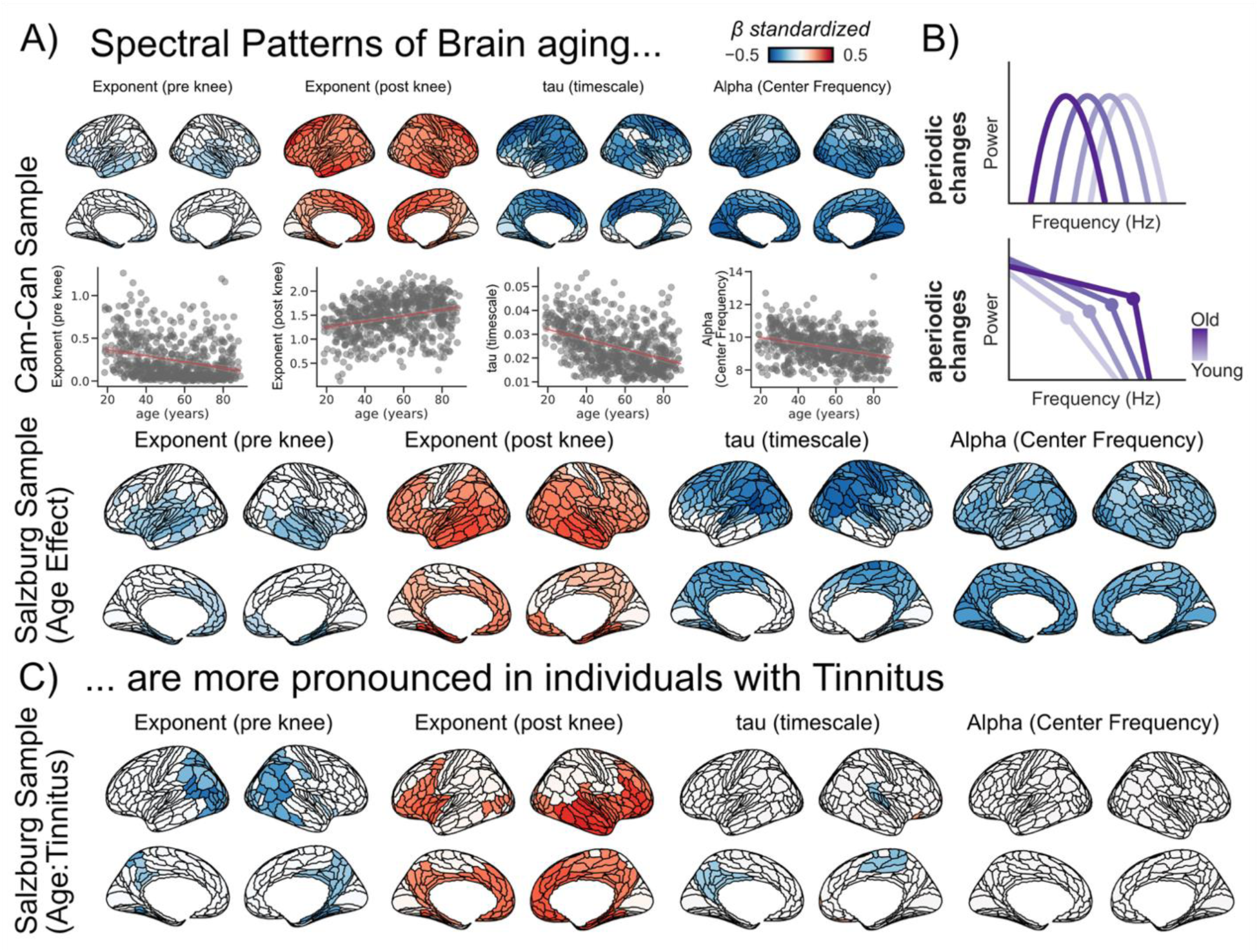
Spectral patterns of brain ageing are more pronounced in individuals with tinnitus. (A) Parcel-wise age effects for key spectral parameters derived from source-level spectral parameterization. Results from the independent Cam-CAN reference sample (top row) replicate and extend canonical ageing signatures, which are also evident in the Salzburg sample (middle row):alpha center frequency slowing, shortening of neural timescales (τ), flattening of the pre-knee exponent, and steepening of the post-knee exponent. (B) Conceptual schematic illustrating how ageing can manifest as periodic changes (e.g., frequency shifts) and aperiodic changes (e.g., slope/exponent and Knee Frequency (i.e. time scale) alterations). (C) Age × tinnitus interaction effects in the Salzburg sample indicate that ageing-related spectral trajectories are more pronounced in individuals with tinnitus, including accelerated alpha slowing, stronger age-related reductions in τ and amplification of aperiodic exponent effects (flatter pre- and steeper post-knee exponents with age). Red colors denote increases, and blue colors denote decreases in the parameter with age (or stronger age effects in tinnitus); brain maps show posterior mean standardized β where the 89% HDI does not overlap with a region of practical equivalence.

**Figure 3:**
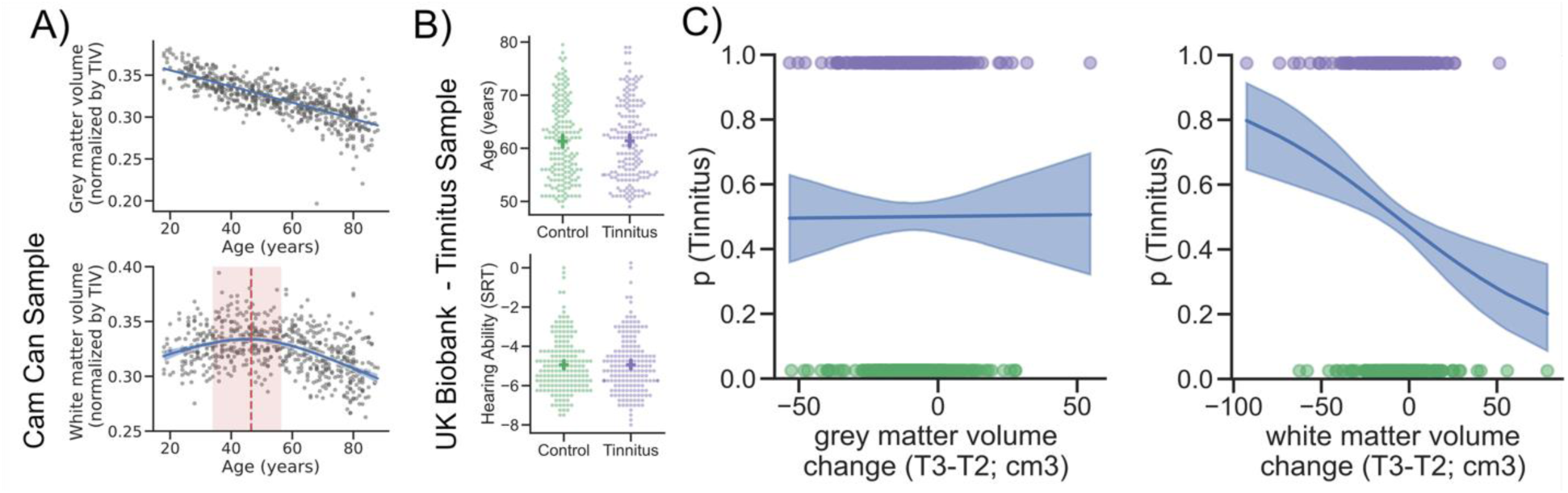
Prospective white-matter decline predicts future tinnitus onset. A) In the Cam-CAN reference sample, total grey-matter volume declines linearly with age, whereas total white-matter volume shows a non-linear age trajectory, increasing in younger adulthood until 46 years (34 – 56 years HDI89%) before decreasing in older adulthood (red dashed line indicates the split). (B) UK Biobank longitudinal sample shows a close to perfect matching based on age and hearing ability (SRT) for controls and individuals with tinnitus. C) Logistic regression models relating longitudinal grey- and white-matter volume changes (T3–T2) to tinnitus probability (T3). Grey-matter change shows no association with tinnitus, whereas greater white-matter volume loss is associated with a higher probability of tinnitus onset. Shaded bands denote 89%HDIs.

To assess whether tinnitus reflects aberrant neural aging, we examined age relationships for both the tinnitus and control groups across all age-sensitive parameters identified in the Cam-CAN reference sample. The results of this analysis show a similar trajectory of age-related spectral changes across tinnitus and controls (see **Figure 2A**; Salzburg sample). Alpha peak frequency exhibited robust age-related slowing (*β_standardized_* = −0.24; range across parcels: −0.39 to −0.15). Similarly, neural timescales decreased (*β_standardized_* = −0.27; range: −0.46 to −0.13), the pre-knee exponent flattened with age (*β_standardized_* = −0.17; range: −0.27 to −0.13), and the post-knee exponent steepened (*β_standardized_*= 0.19; range: 0.10 to 0.31) across both groups.

Notably, all of the aperiodic age effects were amplified in the tinnitus sample (see **Figure 2C**). This effect was most prominent for the aperiodic exponents: the pre-knee exponent showed an enhanced and more widespread age-related flattening (*β_standardized_* = −0.25; range: −0.39 to −0.18), whereas the post-knee exponent showed an enhanced age-related steepening (*β_standardized_* = 0.24; range: 0.19 to 0.36). Also, shortening of neural timescales showed a stronger age-dependency in the tinnitus group, with maximum effects in medial areas (left: posterior medial cortex / precuneus; right: dorsomedial frontal cortex / pre-SMA) and right superior temporal areas (*β_standardized_*= −0.20; range: −0.23 to −0.17). Importantly, as the groups were closely matched for chronological age (**Figure 1A**), the observed effects can therefore not be attributed to age differences per se. Taken together, these cross-sectional MEG findings suggest that age-related trajectories of ongoing neural activity are more extreme and widespread in individuals with tinnitus.

## Prospective white-matter decline predicts future tinnitus onset

The tinnitus-related effects described above were based on cross-sectional comparisons—that is, between individuals who either do or do not report tinnitus at the time of measurement. To more directly test whether aberrant neural aging contributes to tinnitus risk, *longitudinal* data are required from individuals without tinnitus at baseline who later develop chronic tinnitus. However, according to our knowledge, no suitable MEG or EEG datasets of this kind are currently available. Given that neural aging effects are unlikely to be confined to functional measures, we turned to the UK Biobank, which includes repeated structural MRI scans at two time points ([40], separated by approximately 2.73 years (*SD* = 1.33)).

With respect to age-related structural changes, we decided to focus on volumetric changes in white and grey matter. Again, we first turn to the CamCAN dataset to visualize age-related white and grey matter trajectories. On the whole-brain level (**Figure 3A**), we observe the expected decline in grey matter volume with age (*β_standardized_* = -0.76, 89%HDI [-0.80, -0.71]). Other than grey matter, white matter volume exhibited a more complex trajectory. Specifically, white matter increased with age (*β_standardized_* = 0.72, 89%HDI [0.11, 1.29]) up to approximately 46 years (34 – 56 years 89% HDI), followed by a marked decline thereafter (*β_standardized_* = -0.8, 89%HDI [-1.10, -0.49]; see also [41] for converging evidence).

To further assess whether accelerated age-related changes in structural brain differences are predictive of tinnitus onset, we next turned to the longitudinal UK Biobank dataset. From the UK Biobank, we therefore selected individuals who reported new-onset tinnitus at the second assessment (T3) but were tinnitus-free at baseline (T2; *N* = 178). Sex-matched control participants were matched to the tinnitus group for age, and hearing status at T3 (N = 178; **Figure 3C**). Logistic regression analyses revealed that neither grey nor white matter volume at T2 predicted tinnitus onset at T3 (Odds-Ratio_grey matter T2_ = 0.98 89%HDI [0.83, 1.18]; Odds-Ratio_white matter T2_ = 0.95 89%HDI [0.80, 1.11]). Likewise, changes in grey matter volume between T2 and T3 were not predictive of tinnitus onset (Odds-Ratio = 1.00 89%HDI [0.83, 1.17] **Figure 3C**). In contrast, greater white matter volume loss from T2 to T3 was significantly associated with tinnitus emergence at T3 (Odds-Ratio = 0.73 89%HDI [0.61, 0.89] **Figure 3C**). This suggests that structural changes in white, but not grey matter are associated with tinnitus onset. Because hearing status was only matched at T3 between groups, one might still argue that the effect reflects differential hearing decline rather than tinnitus per se. Yet, when we repeated the logistic regression using hearing change (T3–T2) as the predictor, no significant association was observed (Odds-Ratio = 0.99 89%HDI [0.85, 1.21]. Together, these findings indicate that accelerated age-related white matter decline may increase the risk of developing tinnitus, complementing the cross-sectional evidence for aberrant neural aging derived from the MEG data.

## Discussion

The neuroscientific study of tinnitus has largely pursued approaches to identify the “neural correlates“ of this phantom percept. However, this strategy has provided limited leverage on a crucial scientific and clinical question: Why does tinnitus only develop in a subset of individuals exposed to comparable auditory risk factors, such as hearing loss? Here, we advance a novel vulnerability framework in which tinnitus is linked to aberrant trajectories of neural aging rather than being explained solely by the presence of altered auditory activity at a given time (for initial formulations of the idea, see [2]). Two findings anchor this reframing: (i) in cross-sectional resting-state MEG, age–neural relationships in multiple spectral parameters are systematically more pronounced in tinnitus than in closely matched controls, and (ii) in longitudinal UK Biobank imaging, accelerated decline in global white-matter volume predicts subsequent tinnitus onset in participants tinnitus-free at baseline. Crucially, all observed effects remain robust after controlling for major confounders, including chronological age, sex, and hearing status, as well as cardiac activity which we have shown to influence aperiodic activity recorded using M/EEG [31]. Tinnitus distress in our MEG sample was overall very low, providing a very unlikely explanation for our results (see **Supplementary Table 1**). Together, these findings provide population-level, cross-sectional and longitudinal evidence that tinnitus emergence is associated with atypical neural aging trajectories, reframing tinnitus as a disorder of altered brain aging rather than solely aberrant neural activity.

Overall, the neural-aging framework provides a principled explanation for why chronological age is associated with increased tinnitus risk independent of hearing status [2]. Normal aging is characterized by several processes, as also shown in our study, including white-matter decline [41] and alterations in excitatory–inhibitory balance [42], which may disrupt communication across large-scale brain networks. An acceleration or maladaptive expression of these processes in tinnitus could facilitate the emergence of a tinnitus precursor signal [43], for example by allowing activity to escape reduced internal suppression mechanisms [22] and spread beyond auditory processing regions to influence downstream areas [44], or by increasing the precision of priors [43]. Within this framework, tinnitus is not solely a consequence of local auditory dysfunction, but reflects a systems-level imbalance in network dynamics [45]. This perspective may also help explain why the “neural correlates” approach, which has dominated the tinnitus electrophysiology literature to date, has yielded relatively modest and inconsistent findings: different studies may sample distinct points along a heterogeneous aging–plasticity trajectory, and may conflate periodic oscillatory activity with aperiodic background dynamics unless these components are explicitly separated.

The latter aspect—namely the contribution of aperiodic activity—is increasingly recognized as critically important, yet has largely been overlooked in tinnitus research, which has traditionally relied on band-limited power analyses [12]. Within this framework, it is conceivable that commonly reported increases in low-frequency activity (e.g., delta) may, at least in part, reflect a steepening of the (pre-knee) aperiodic slope, as observed here at the group level (see also pre knee Exponent effect in **Supplementary Figure S2**). This interpretation is further supported by the overall low probability of detecting genuine oscillatory peaks in the low-frequency range (**Figures 1B** and **1D**), suggesting that apparent “oscillatory” effects may instead arise from broadband spectral changes. Beyond clarifying low-frequency findings, spectral parametrization enables a more precise characterization of alpha activity in tinnitus. While alpha remains the most prominent rhythmic component also in tinnitus, we observe a pronounced reduction—often manifesting as an absence—of alpha activity in left auditory cortex, extending our earlier report of reduced alpha power in a similar region obtained without parametrization [14]. This effect is complemented by reduced alpha peak height in left sensorimotor regions. Alpha oscillations have long been linked to inhibitory gating [34, 46], a view supported by invasive animal work demonstrating rhythmic suppression of spiking with stronger alpha [47], as well as human studies linking alpha to perceptual reports such as in near threshold tasks [16]. Importantly, reduced alpha does not appear to enhance perceptual sensitivity per se, but rather shifts perceptual decision criteria toward a more liberal regime [48]. Taken together, these findings support the notion that reduced alpha activity in tinnitus may reflect diminished rhythmic inhibitory control, thereby facilitating the propagation of internally generated auditory activity [15, 49, 50].

Beyond replicating and refining alpha power effects, the parametrization approach also revealed novel alterations not captured by conventional band-limited analyses. Specifically, we observed increased alpha peak frequency in bilateral frontal cortices and reduced alpha bandwidth in posterior cingulate regions encompassing the precuneus. Alpha peak frequency has been proposed to reflect the temporal resolution or “sampling rate” of cortical processing [51], with converging evidence linking it to inhibitory neurochemistry, including GABA levels [52] and receptor kinetics [53]. Notably, most prior work has focused on posterior regions, particularly occipital cortex; the present frontal effects therefore suggest that tinnitus-related alterations extend to higher-order control systems, potentially reflecting changes in the regulation of internally generated representations. In parallel, the reduced alpha bandwidth in precuneus indicates a more spectrally constrained and less variable oscillatory regime in a key hub of the default mode network. Given the established role of this region in internally oriented cognition and conscious awareness [54], as well as prior evidence linking it to tinnitus-related functional connectivity and distress [55, 56], this finding points to altered intrinsic dynamics in large-scale networks supporting internal mentation. Elucidating the circuit-level mechanisms underlying such bandwidth changes remains an important avenue for future work.

A major advantage of the present approach is the ability to characterize aperiodic activity, which has been largely absent from prior tinnitus research. We observed robust steepening of both pre- and post-knee aperiodic slopes, with distinct spatial distributions: pre-knee effects were primarily localized to posterior and anterior midline regions, whereas post-knee effects were mainly right-lateralized across temporal, parietal, and sensorimotor cortices. The aperiodic exponent has been widely interpreted as a proxy for excitatory–inhibitory (E/I) balance [37], with steeper slopes suggesting relatively increased inhibitory influence. This interpretation is supported not only by computational modeling, but also by empirical work linking the exponent to arousal states [57, 58] and pharmacological manipulations [59]. Leveraging the model proposed by Gao et al. [37], which simulates local field potentials as a combination of excitatory (AMPA) and inhibitory (GABA) synaptic inputs, it becomes apparent that E/I-related effects are strongly reflected in the post-knee exponent, and related parameters such as the knee frequency and neural timescale (τ) (see **Supplementary Figure S3**). Notably, we did not observe consistent group differences in these latter parameters, arguing against a straightforward interpretation in terms of globally altered E/I balance (see also [60, 61]). Future work integrating noninvasive human recordings with invasive tinnitus animal models will be critical for establishing the underlying circuit mechanisms driving the aperiodic group differences.

Taken together, the findings discussed above indicate that tinnitus is characterized by a multi-faceted reorganization of cortical dynamics, encompassing oscillatory, aperiodic, and spatially distributed alterations that are not readily captured by traditional band-limited approaches. However, a central premise motivating the present work is that these alterations may not simply reflect static disease markers but rather emerge from aberrant trajectories of neural aging that increase vulnerability to tinnitus. To address this, we first established normative age relationships using an independent lifespan dataset [30], and observed closely matching patterns in our own data. These included previously reported age-related changes such as decreased alpha frequency [38, 39], a flattening of the pre-knee and steepening of the post-knee aperiodic slope [31, 42], and shortened neuronal timescales [62, 63]. Crucially, for all aperiodic parameters—but not for alpha peak frequency—these age-related effects were systematically amplified in the tinnitus group. This suggests that tinnitus is associated with a shift toward a more extreme position along aging-sensitive dimensions of cortical dynamics. Computational modeling (see **Supplementary Figure S3**) indicates that this parameter is most closely linked to the relative dominance of excitation over inhibition, providing a mechanistic bridge between macroscopic spectral features and underlying circuit dynamics. Hence, the shortening of neuronal timescales appears particularly informative for interpreting the underlying circuit-level changes. Spatially, these effects encompass medial sensorimotor regions, the precuneus, and right auditory cortex, pointing to a distributed network-level alteration. This pattern is consistent with the notion that aging is accompanied by a decline in inhibitory efficacy, particularly in GABAergic systems, which in the context of tinnitus may interact with already reduced rhythmic inhibitory control (as reflected in alpha alterations) or a diminished capacity to suppress internally generated noise. In this light, the observed spectral alterations are unlikely to represent a singular “neural correlate” of tinnitus. Rather, they point to a systems-level vulnerability that arises from an interaction between aging-related changes in cortical dynamics and additional risk factors such as peripheral hearing damage. This perspective reframes tinnitus not as a localized dysfunction, but as an emergent consequence of altered large-scale brain dynamics unfolding along an accelerated or dysregulated aging trajectory. To move beyond cross-sectional inference and directly address vulnerability, longitudinal approaches are required to determine whether these neural signatures precede and predict tinnitus onset.

Longitudinal M/EEG datasets of sufficient scale to directly test these ideas are currently lacking. However, well-characterized aging trajectories also exist for structural brain features [64], including global reductions in grey and white matter, as observed here in the CamCAN dataset (see **Figure 3**). If aberrant neural aging represents a general vulnerability principle for tinnitus—rather than being restricted to dynamic activity alone—then more pronounced age-related structural decline should be associated with an increased likelihood of developing tinnitus. To test this prediction, we leveraged a large prospective sample from the UK Biobank, focusing on individuals without tinnitus at baseline. We examined whether baseline global grey and white matter density, as well as longitudinal changes in these measures, were associated with the emergence of tinnitus at follow-up. Strikingly, we found that greater reductions in white matter were associated with an increased risk of subsequently reporting chronic tinnitus. This effect could not be explained by baseline variables such as age, sex, or hearing loss, nor by changes in hearing thresholds over time, due to careful matching and analysis procedures. To our knowledge, this represents the first identification of a prospective neural risk marker for tinnitus and provides independent support for the proposed aberrant aging framework. White matter integrity is critical for efficient long-range communication and the coordination of distributed neural activity [65]. Its decline, therefore, may reflect a reduced capacity to constrain and regulate internally generated activity across large-scale networks. In this context, greater white matter loss may render the brain more permissive to the emergence and persistence of internally generated auditory signals. This interpretation aligns with the functional MEG findings, particularly the steeper post-knee aperiodic slope observed in frontal and parietal regions (**Figure 2C**), which has been linked to alterations in excitatory–inhibitory balance. While the precise relationship between these structural and functional changes remains to be established, their convergence is consistent with a model in which tinnitus arises from an interaction between local circuit alterations and a breakdown of large-scale network regulation along aging-sensitive trajectories. Taken together, these findings suggest that white-matter decline constitutes a prospective vulnerability factor for tinnitus, supporting the notion that altered trajectories of brain aging increase the likelihood of developing persistent phantom percepts.

Despite representing a significant advance, the present study has several important limitations. Most critically, neither the cross-sectional nor the prospective analyses allow for causal inference. It remains conceivable that the observed neural aging signatures arise as a consequence of tinnitus, or reflect epiphenomenal processes rather than predisposing factors. Addressing this question will require a closer integration with animal models of tinnitus, in which aging-related processes can be experimentally manipulated—for example through genetic or environmental interventions [66]—prior to inducing auditory damage. Such approaches would enable direct testing of the central prediction of our framework, namely that alterations in neural aging trajectories modulate vulnerability to the development of tinnitus-like behavior. A second limitation concerns the still incomplete mechanistic understanding of the spectral measures derived from parametrization approaches. While these measures provide a powerful decomposition of neural activity, their precise physiological underpinnings remain an active area of research. Progress in this domain will depend on continued integration of computational modeling, pharmacological manipulations, and invasive recordings [60]. Finally, the relationship between the functional MEG findings and the structural MRI results remains indirect. Establishing a mechanistic link between these levels of description will require large-scale longitudinal studies that combine electrophysiological and structural measures within individuals over time.

Overall, our findings have potential implications for treatment strategies targeting both rhythmic inhibitory control and broadband activity putatively linked to excitatory–inhibitory (E/I) balance. Neurofeedback approaches have shown promising initial effects [67], but remain limited by practical constraints (e.g., prolonged training, scalability) and methodological challenges, including the signal-to-noise limitations of EEG. More broadly, the present results support a shift away from a narrow search for “neural correlates” of tinnitus perception toward a framework in which tinnitus reflects an underlying vulnerability emerging from altered neural aging. In this view, changes in connectivity, E/I balance, and intrinsic dynamics collectively increase the likelihood that internally generated auditory activity is experienced as a persistent percept [43]. This perspective opens new avenues for both intervention and prevention. Future work will need to determine to what extent these neural signatures can be leveraged for individual risk profiling and early identification of vulnerable individuals. Importantly, our findings also motivate a broader preventive approach that extends beyond hearing loss and targets aging-related neural processes themselves. From this perspective, progress in tinnitus research may critically depend on closer alignment with advances in the study of healthy brain aging.

## Methods

### Sample

The present study builds on electrophysiological (MEG) resting-state data and structural recordings (MRI) from several sources. The MEG data analyzed in this study was obtained from two different sources: the CamCAN repository [30] and resting-state data routinely recorded at the University of Salzburg. The sample obtained from CamCAN contained 655 healthy volunteers with an age range between 18 and 88 years of age with an average age of 54 and a standard deviation of 18 years. The sample obtained from the University of Salzburg contained 106 volunteers with an age range between 20 and 76 years of age with an average age of 53 and a standard deviation of 15 years. 53 of the subjects measured at the University of Salzburg reported that they experience tinnitus. Among participants who reported tinnitus, tinnitus distress was generally low, as measured using the Mini-TQ12 [68], with 79% reporting low distress (see **Supplementary Figure S1**).

The other 53 subjects were selected from the pool of resting state measurements, routinely recorded at the University of Salzburg, and were matched based on age, hearing ability and sex. Tinnitus, sex and age were assessed using an ongoing online study of hearing epidemiology in the county of Salzburg (Austria) (SalzburgHört; https://salzburghoert.sbg.ac.at/). Hearing ability was measured using standard pure tone audiometry with the AS608 Basic (Interacoustics, Middelfart, Denmark) and the pure tone average across both ears and all frequencies (125 - 8000Hz) was computed. Afterwards we queried our database of resting-state MEG recordings to find a subject that was optimally matched in hearing ability, age and sex to a tinnitus subject (see also **Figure 1A**). The 53 tinnitus subjects reflect the full sample of subjects with tinnitus at the time of querying our database for the present analyses (2023). The MRI data analyzed in this study was obtained from two different sources: the CamCAN repository [30] and the UK Biobank (https://www.ukbiobank.ac.uk/). The CamCAN sample used for the MRI analyses was equivalent to the MEG CamCAN sample described above. The UK Biobank is a major biomedical database comprising a total of 502,382 participants which provided questionnaire data, physical tests, genetic information, and imaging data. The data of the UK Biobank was recorded in four collection periods (T0, T1, T2 & T3) and MRI data was available only for T2 and T3. From the UK Biobank population, we selected a matched group of subjects that were similar in age, sex and hearing ability (at T2 & T3), but differed in either having or not having tinnitus at T3 (see also **Figure 3C**). As an additional constraint, all selected subjects had to be tinnitus free at T2 and never reported tinnitus in the past (T0 and T1). Finally, we only selected data from subjects that did not report a history of psychiatric care on admission and only selected those where no incidental findings were made during any of the MRI scans. This resulted in a total of 356 subjects with an age range between 49 and 79 years of age with an average age of 61 and a standard deviation of 7 years. 178 of the subjects reported that they experience tinnitus at T3. Data was only excluded from further analysis when the automatic preprocessing procedure (see *MEG preprocessing & MRI preprocessing*) resulted in errors.

The CamCAN study was conducted in compliance with the Helsinki Declaration and has been approved by the local ethics committee, Cambridgeshire 2 Research Ethics Committee (reference: 10/H0308/50). Ethical approval for UK Biobank was gained from the Research Ethics Service (REC reference: 21/NW/0157), and written informed consent was obtained from all participants. This project was accepted under the project ID 884663. The MEG data obtained at the University of Salzburg were recorded as part of routine resting-state measurements conducted prior to various experimental paradigms, which were approved by the Ethics Committee of the University of Salzburg. Informed consent was obtained from each participant.

## MEG processing

### Data Acquisition

MEG data (CamCAN) was recorded at the University of Cambridge, using a 306 VectorView system (Elekta Neuromag, Helsinki). MEG data (Salzburg) was recorded at the University of Salzburg, using a 306 channel TRIUX system (MEGIN Oy, Helsinki). Both systems are equipped with 102 magnetometers and 204 planar gradiometers and positioned in magnetically shielded rooms. In order to facilitate offline artifact correction, electrooculogram (VEOG, HEOG) as well as ECG were measured continuously in both recording sites. Data recorded at Cambridge was online filtered at 0.03–330 Hz, whereas data recorded at Salzburg was online filtered at 0.1–330 Hz with a 1000 Hz sampling rate at both recording sites. Five Head-Position Indicator coils were used to measure the position of the head. All data used in the present study contains passive resting-state measurements lasting about ∼8 min (Cambridge) and ∼5 min (Salzburg). Further data processing at both recording sites was conducted similarly and will therefore be reported together.

### MEG preprocessing

Data preprocessing was performed using Python (Version 3.12.8) and MNE Python (Version 1.8). In the first step, a signal space separation algorithm was used to clean the data from external noise. All data were high pass filtered at 0.1 Hz (Kaiser windowed finite impulse response filter). To identify eye-blinks and cardiac components in the data, 50 independent components were calculated using the *fastica* algorithm ([69]; implemented in MNE-Python *version 1.2*; with the parallel/symmetric setting; note: 50 components were selected for MEG for computational reasons). As ICA is sensitive to low-frequency drifts, independent components were calculated on a copy of the data high-pass filtered at 1 Hz. Components related to cardiac and ocular activity were determined via correlation with concurrent ECG, VEOG, and HEOG recordings. A threshold of *r*>0.4 was applied to detect the ECG components and a threshold *r*>0.4 for detecting EOG components in the data and components in which the correlation exceeded said threshold were removed.

Afterwards, the data were projected to source space, to better understand the neural sources contributing to the signal measured at the MEG sensors. A semi-automatic pipeline was used to co-register the individual subjects’ head shape with a "standard" model created by the combination of 40 MRI scans of real brains (*fsaverage*; [70]). This approach has been shown to lead to comparable results as compared to manual co-registration [71]. The co-registered head models were then used to calculate the forward solution. First, we computed the single-layer boundary element model (BEM; [72]) to create a BEM solution for the *fsaverage* template brain. Second, to define the position and orientation of the sources, we create a decimated dipole grid on the grey matter surface using an icosahedron subdivision (ico-4) which includes 2562 sources for hemispheres. Assuming a cortical surface of 1000 cm^2^, the approximate spacing between the grid points was 6.2 mm, and each voxel occupied 39 mm^2^ of the cortical surface. To control for field patterns related to noisy sources (e.g., human-based or environmental), we weighed each channel by a noise covariance matrix computed on the nearest empty room recording. That is, prior to any daily data acquisition, ∼2 min of empty room measurements (data collected with no subject) were recorded in the MEG; depending on the date of testing, the closest measurement was found for each subject and preprocessed in the exact same way [73]. The Noise covariance matrix and the true rank were computed from the empty room measurement and used together with the data covariance matrix to calculate the inverse solution based on common linearly constrained minimum variance (LCMV) beamformer spatial filters [74]. These spatial filters were then applied on the preprocessed MEG data to project the signal to source space. Afterwards the source projected data were parcellated using the HCP-MMP1 atlas [75] and the average across vertices for each time-point within each label of the atlas was computed, by finding the dominant direction of source space normal vector orientations within each label of the atlas and applying a sign-flip to time series at vertices whose orientation is more than 90° different from the dominant direction.

### Spectral analysis

Power spectra were computed using Welch’s method [76] between 1 and 100 Hz (0.25 Hz resolution) on the parcellated timeseries data. Aperiodic activity was extracted using the IRASA algorithm [29] implemented in the PyRASA package [28]. Afterwards, periodic and aperiodic components were identified. Periodic components of narrowband neural activity were extracted from the “periodic spectrum” (returned by subtracting the aperiodic signal obtained by via IRASA from the original welch power spectrum) via a peak finding procedure implemented in PyRASA based on the “find_peaks” algorithm implemented in SciPy [77]. This resulted in periodic components (Power, Center Frequency and Bandwidth) for each of the canonical frequency bands (Delta (1-3Hz), Theta (3-7Hz), Alpha (7-14Hz) & Beta (15-30Hz); selected canonical frequency band definitions are based on [27]). The aperiodic parameters were identified via fitting the following model to the aperiodic spectrum obtained from IRASA.

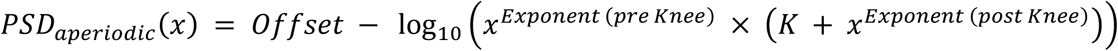

We opted for modelling the data by fitting two different exponents (pre and post Knee) as opposed to the Lorentzian function proposed in [27] that models only the Knee and an Exponent after the Knee. This modelling decision was taken, because we noticed that both the estimation of the Knee and Exponent (and thereby also the timescale estimate tau) was significantly off, whenever the slope of the spectrum differed from 0 before a Knee or bend in the spectrum was observable (which is a common sight in MEG and EEG recordings). Allowing for two slopes mitigated this problem both in simulated and observed data.

### MRI processing

#### MRI preprocessing

Structural MRI data from the CamCAN repository were processed using FastSurfer [78], which generates FreeSurfer-compatible cortical surface reconstructions and anatomical segmentations. For each subject, T1-weighted (T1w) and T2-weighted (T2w) images were provided as inputs to the FastSurfer pipeline (run_fastsurfer.sh). Hypothalamus processing was disabled (--no_hypothal). The resulting subject-specific FreeSurfer directories were subsequently used for atlas-based parcellation and region-wise morphometric analyses. To obtain HCP-MMP1 atlas labels in each subject’s native surface space, the HCP-MMP1 annotation files were transferred from fsaverage to each subject using FreeSurfer’s mri_surf2surf [70, 79]. These subject-level HCP-MMP1 labels were then used to derive parcel-wise morphometric summaries (including volumetric measures) for downstream statistical analyses. For UK Biobank participants, we used the structural MRI-derived measures provided by UK Biobank (generated by their standard imaging processing pipeline) and did not apply additional structural preprocessing.

### Statistical Inference

To investigate the relationship between tinnitus, age and structural (MRI) or functional (MEG) differences we used bayesian generalized linear models (GLMs) built in PyMC (a python package for probabilistic programming; [80]). The model structure used for inference was based on simulations (see *Simulations*) and is described below (*Statistical Models*). Priors were chosen to be weakly informative thereby offering regularization for unrealistically large effects.

#### Statistical Model MEG

Choosing a good statistical model for MEG data is not straightforward, as it depends, among other things, on our assumptions for the spatial distribution of the data generating process behind the modeled data. Here we assume that A) local brain activity is nested in larger cortical areas and that B) larger cortical areas can be independently modulated. This means that while we expect that activity in different parcels in a cortical area is similarly modulated, we do not explicitly assume the presence of whole brain effects. We therefore model the effect of Age, Tinnitus and their interaction on spectral changes in the MEG using the generalized linear mixed effect model described below.

Let:

- *i* = 1, …,*N* : observations
- *parcel =* 1, …,*R : parcels* (HCP-MMP1 parcellation)
- cortex = 1, …, C : cortices (HCP-MMP1 parcellation)
- *parcel(i)* : parcel of an observation *i*
- *cortex(parcel) :* cortex in which a parcel is located
- *k* indices of the effect labels (Age, Tinnitus, Age x Tinnitus)
- *y* = the modeled outcome (in the case of MEG a spectral feature (e.g. alpha center frequency)

Model:

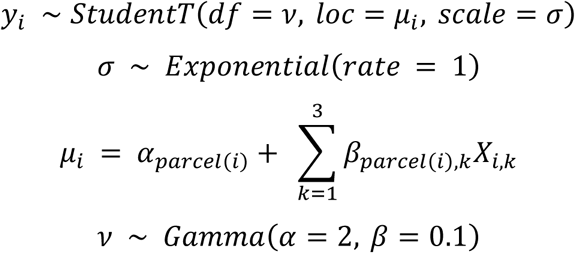

Design matrix:

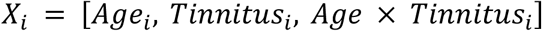

parcel-level priors:

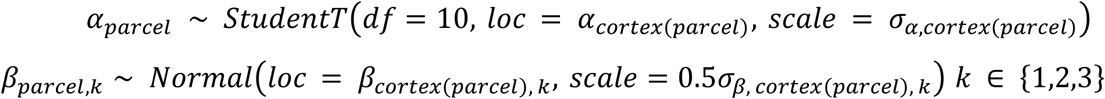

Cortex-level priors:

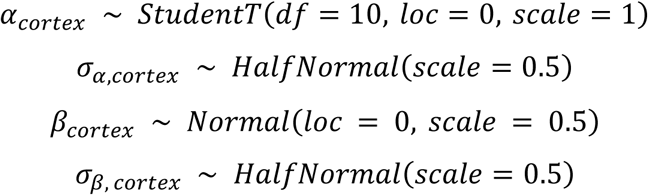

For binary outcomes (e.g. the presence of an alpha oscillation), we used a Bernoulli likelihood with a logit link.

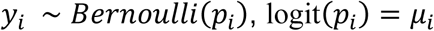

Before statistical inference, the continuous predictor age and continuous neural outcome variables (e.g., alpha center frequency) were z-scored. Tinnitus status was effect-coded (absence = −0.5, presence = +0.5) to facilitate interpretation of main effects in the presence of an interaction. With this coding, the tinnitus main effect reflects the difference between participants with and without tinnitus at mean age (age z = 0), which is sensible given the wide age range covered in our study. The main effect of age reflects the average association between age and the neural outcome across tinnitus groups (i.e., at tinnitus = 0). The interaction term reflects the difference in the age–outcome slope between participants with and without tinnitus.

#### Simulations MEG Model

To evaluate the validity of our statistical model, we simulated data under the assumptions specified above (see Statistical Model and **Supplementary Figure S4B**) and compared its performance to two commonly used alternatives with similar prior specifications. Model #1 estimated parcel-specific effects without pooling (**Supplementary Figure S4A**; Unpooled Model). Model #2 assumed a single hierarchical level across all parcels, yielding partial pooling toward a global (whole-brain) effect (**Supplementary Figure S4A**; LME-Model). Model #3 assumed that parcels were nested within cortical areas, yielding partial pooling toward cortex-specific effects (**Supplementary Figure S4A**; Cortex-LME-Model; see Statistical Model).

The simulation aimed to assess which model best recovered the simulated effects while maintaining a low false-positive rate. For each simulation run (N_runs_ = 100), we generated data for *N*=40 subjects and all parcels using a standardized subject-level predictor *X* (z-scored) and simulated parcel-wise outcomes *y* as a linear function of *X* with additive Gaussian noise (SD = 1). parcels belonging to a cortex of interest (e.g., Primary Auditory cortex) were assigned a non-zero slope (randomly sampled from a predefined set of values between 0.3 and 0.8), whereas all other parcels were simulated under the null (slope = 0; see also **Supplementary Figure S4B**). Afterwards, the three models were evaluated for their ability to correctly identify simulated effects while avoiding spurious effect detections in parcels simulated under the null (**Supplementary Figure S4C**). Overall, the Cortex-LME-Model provided the best trade-off, achieving the highest true-positive detection rate while maintaining a low false-positive detection rate comparable to the whole-brain LME-Model. Although the whole-brain LME-Model yielded the lowest false-positive rate, it also failed to recover most simulated effects. We additionally assessed interval calibration by computing, across simulation runs, whether the true simulated effect size was contained within the 89% highest density interval (HDI) of the posterior slope estimate (coverage). Consistent with the detection results, the Cortex-LME-Model showed the most favorable coverage for non-null effects across parcels while retaining good coverage under the null (see **Supplementary Figure S4D**). In sum, these simulations support the use of cortex-level partial pooling when (A) local activity is structured by larger cortical areas and (B) cortical areas can vary independently.

## Statistical Model MRI

The age-related changes in MRI data were analyzed on whole brain level. In the CamCAN dataset the volume reflects the whole brain white and grey matter volumes normalized by the total intracranial volume (TIV). In the UKBiobank data, the white and grey matter volumes were normalized using a head size scaling factor. Both the volumetric data and the age of the participants were z-scored before subjecting them to a statistical model. Considering the linear decline of grey matter with age grey matter decline was modeled using a simple linear regression model with the following priors.

Let:

*y* = Grey Matter Volume (normalized by TIV) and standardized across subjects

*X* = Age (standardized across subjects)

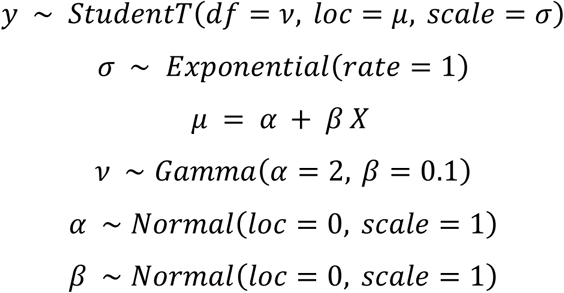

A similar model structure and prior setup was used when using the change in grey/white matter to predict tinnitus onset with some slight changes. Here *X* reflects the grey/white matter change and *y* encodes whether someone has tinnitus or not. Consequently, Instead of the StudentT likelihood to model *y* a Bernoulli likelihood with a logit link was used resulting in the following change to the model structure.

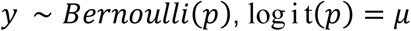

When modeling the age-related change in overall white matter volume, we noticed that the change of white matter volume was non-linear, with an age-related increase in younger subjects followed by a decrease in older individuals (see **Figure 3A**). We wanted to catch this non-linearity and its changepoint and therefore modeled the data as such:

Let:

*y* = White Matter Volume (normalized by TIV) and standardized across subjects

*X* = Age (standardized across subjects)

*CP* = Changepoint centered around 45 years and standardized across subjects

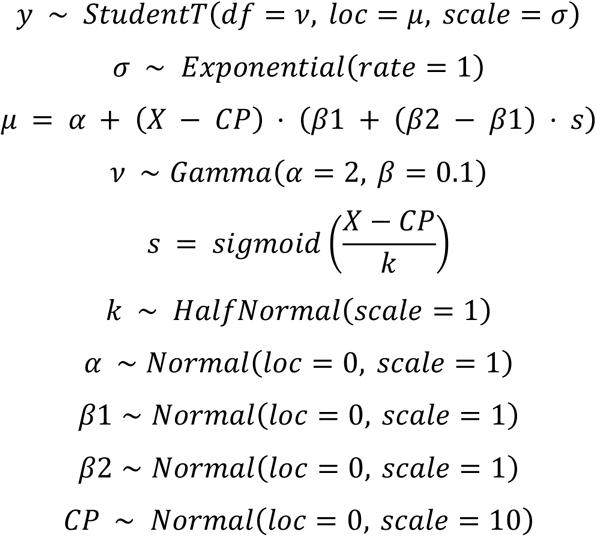

## Statistical Significance

Results were considered statistically significant if 89% of the highest (probability) density interval (HDI) of the posterior for a given standardized β-coefficient was not overlapping with a region of practical equivalence between -0.05 and 0.05. We chose this less critical criterion as the more common –0.1 and 0.1 threshold based on small effect sizes by [81], as our simulations have shown that with a less conservative bound, we were more likely to recover “true” simulated effects in the range of 0.3 to 0.7 (which reflect sensible effect sizes of interest in M/EEG data analysis) compared to the more conservative bound, while retaining a very low error rate (< 0.1%). Furthermore, it was ensured that for all models, there were no divergent transitions (R_hat_ < 1.05 for all relevant parameters) and an effective sample size > 400 (an exhaustive summary of bayesian model diagnostics can be found in [82]).

## Data visualization

Individual plots were generated in Python using Matplotlib [83] and Seaborn. [84]. Parcellated brain plots were plotted using custom plotting functions (https://github.com/schmidtfa/ggseg_py.git) that take advantage of GeoPandas [85] and ggseg [86]. Plots were then arranged as cohesive figures with Affinity Designer (https://affinity.serif.com/en-us/designer/).

## Acknowledgements

This project received financial support from the Land Salzburg through the BrainAge project (20102/F2400537-FPR). This research has been conducted using the UK Biobank Resource under application number 884663. Data collection and sharing for this project was provided by the Cambridge Centre for Ageing and Neuroscience (CamCAN). CamCAN funding was provided by the UK Biotechnology and Biological Sciences Research Council (grant number BB/H008217/1), together with support from the UK Medical Research Council and University of Cambridge, UK. The authors also acknowledge the computational resources and services provided by Salzburg Collaborative Computing (SCC), funded by the Federal Ministry of Education, Science and Research (BMBWF) and the State of Salzburg. We furthermore want to thank Lisa Reisinger and Manfred Seifter for their support in data collection. ChatGPT-5 was used for language editing to improve readability and clarity. The authors reviewed and approved all final wording.

## Author contributions

Conceptualization: NW, FS Formal Analysis: FS, NW Investigation: NW, FS Methodology: FS, NW Supervision: NW Visualization: FS

Writing—original draft: NW, FS

Writing—review & editing: NW, FS, GD, NMV, SMK, ET

## Data & Code availability

The MEG data obtained at the University of Salzburg will be made publicly available on the https://anc.plus.ac.at/ upon publication. The MEG/MRI data used for the healthy ageing analyses are obtained from https://cam-can.mrc-cbu.cam.ac.uk/functional/. The MRI data used in the prospective tinnitus analysis was obtained from the UK Biobank. All code used for the analysis is publicly available on GitHub at: https://github.com/schmidtfa/resting_tinnitus

## Supplementary Material

Tinnitus Distress (Mini-*TQ12*)

**Table 1:**
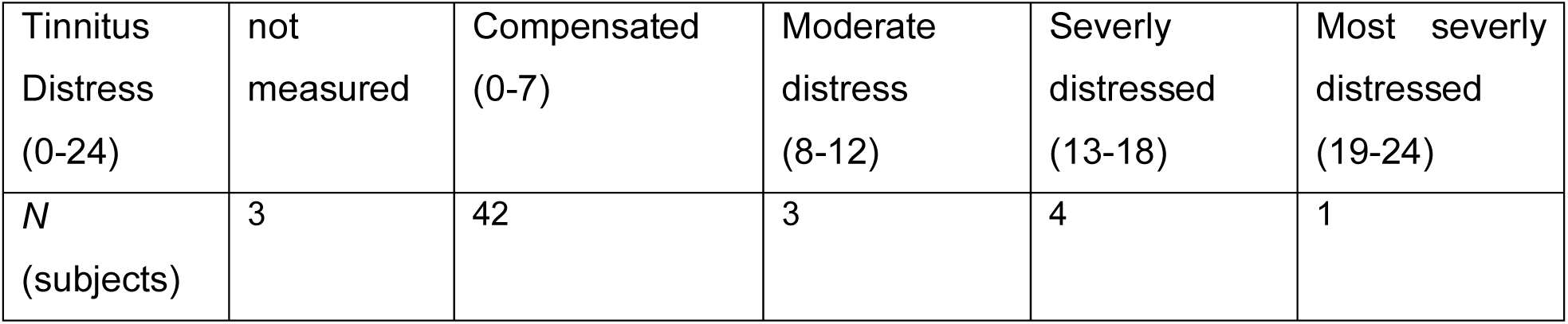
Tinnitus Distress across individuals with tinnitus.

**Table 2:** Model summary statistics Exponent (pre Knee)

|  | B (standardized) | SD | HDI 5.5% | HDI 94.5% |
| --- | --- | --- | --- | --- |
| Intercept | 0.380 | 0.006 | 0.371 | 0.389 |
| Age | 0.008 | 0.007 | -0.003 | 0.018 |
| Tinnitus | 0.009 | 0.012 | -0.007 | 0.026 |
| Age:Tinnitus | -0.006 | 0.017 | -0.027 | 0.012 |

Aperiodic findings are not significantly confounded by cardiac artifacts

**Table 3:**
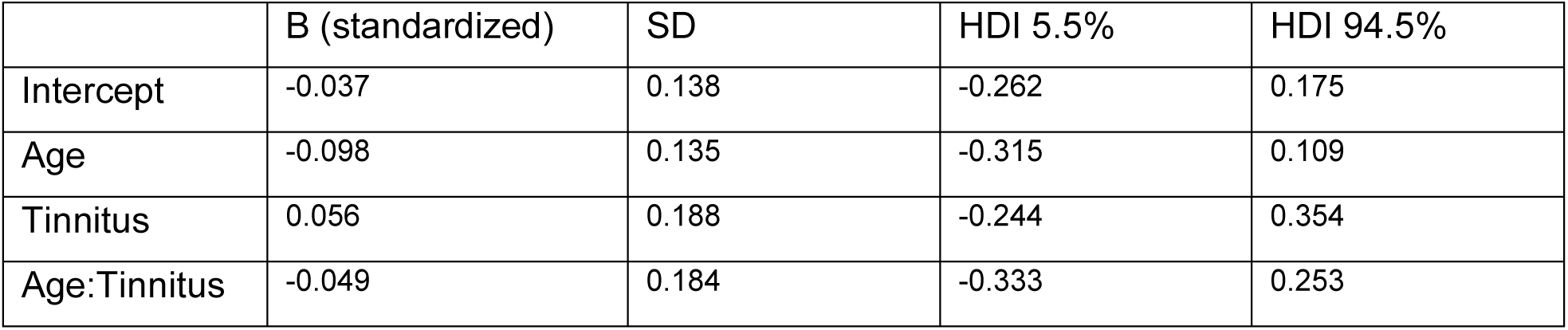
Model summary statistics Exponent (post Knee)

|  | B (standardized) | SD | HDI 5.5% | HDI 94.5% |
| --- | --- | --- | --- | --- |
| Intercept | -0.037 | 0.138 | -0.262 | 0.175 |
| Age | -0.098 | 0.135 | -0.315 | 0.109 |
| Tinnitus | 0.056 | 0.188 | -0.244 | 0.354 |
| Age:Tinnitus | -0.049 | 0.184 | -0.333 | 0.253 |

**Supplementary Figure S1.**
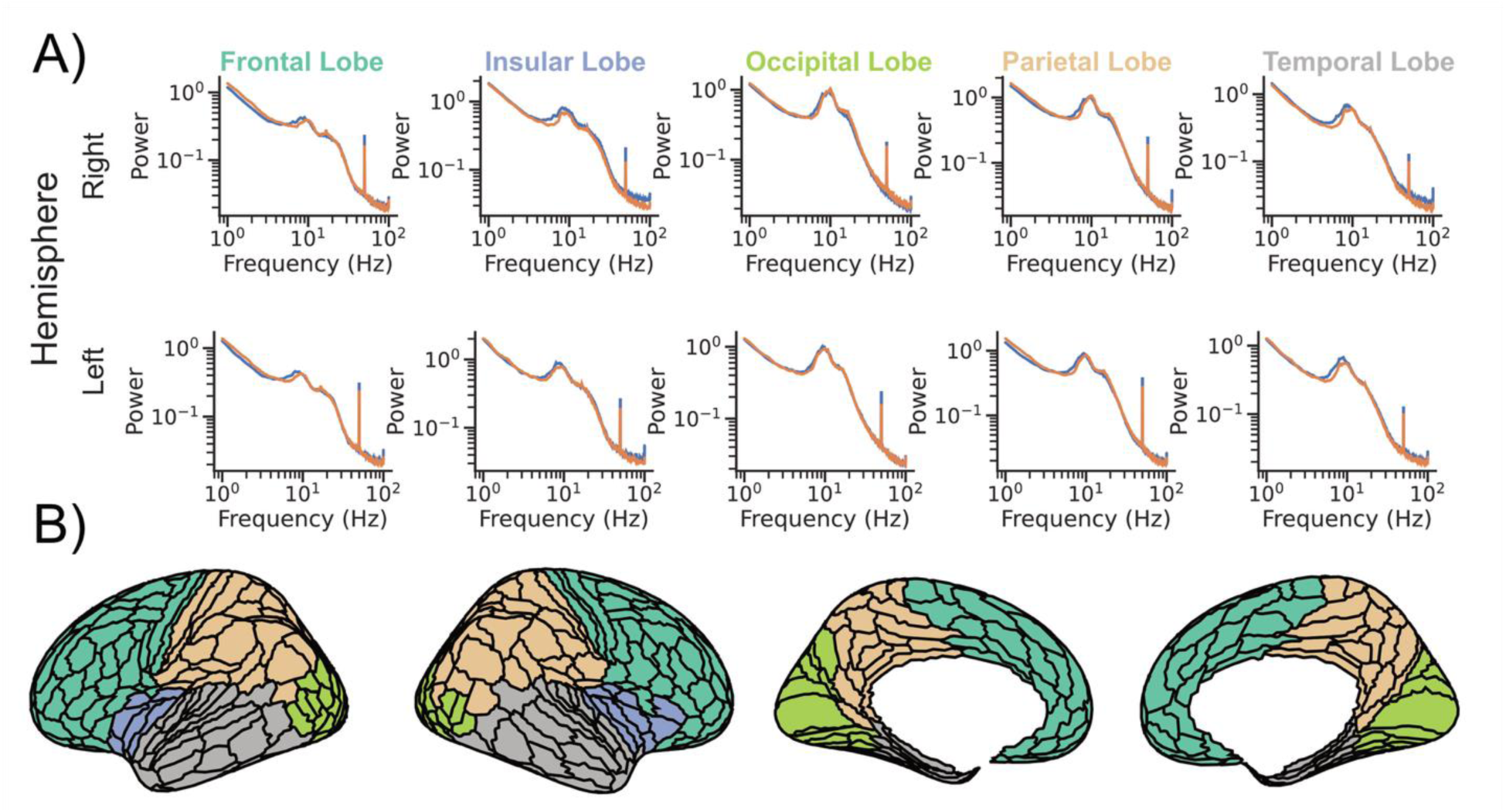
: Lobar distribution of raw power spectra across the cortex. **A)** Group-averaged raw power spectra for participants with tinnitus (orange) and controls without tinnitus (blue), shown separately by hemisphere and across the five lobes of the HCP-MMP1 parcellation. **B)** Cortical surface renderings illustrating the lobe assignments of HCP-MMP1 regions.

**Supplementary Figure S2.**
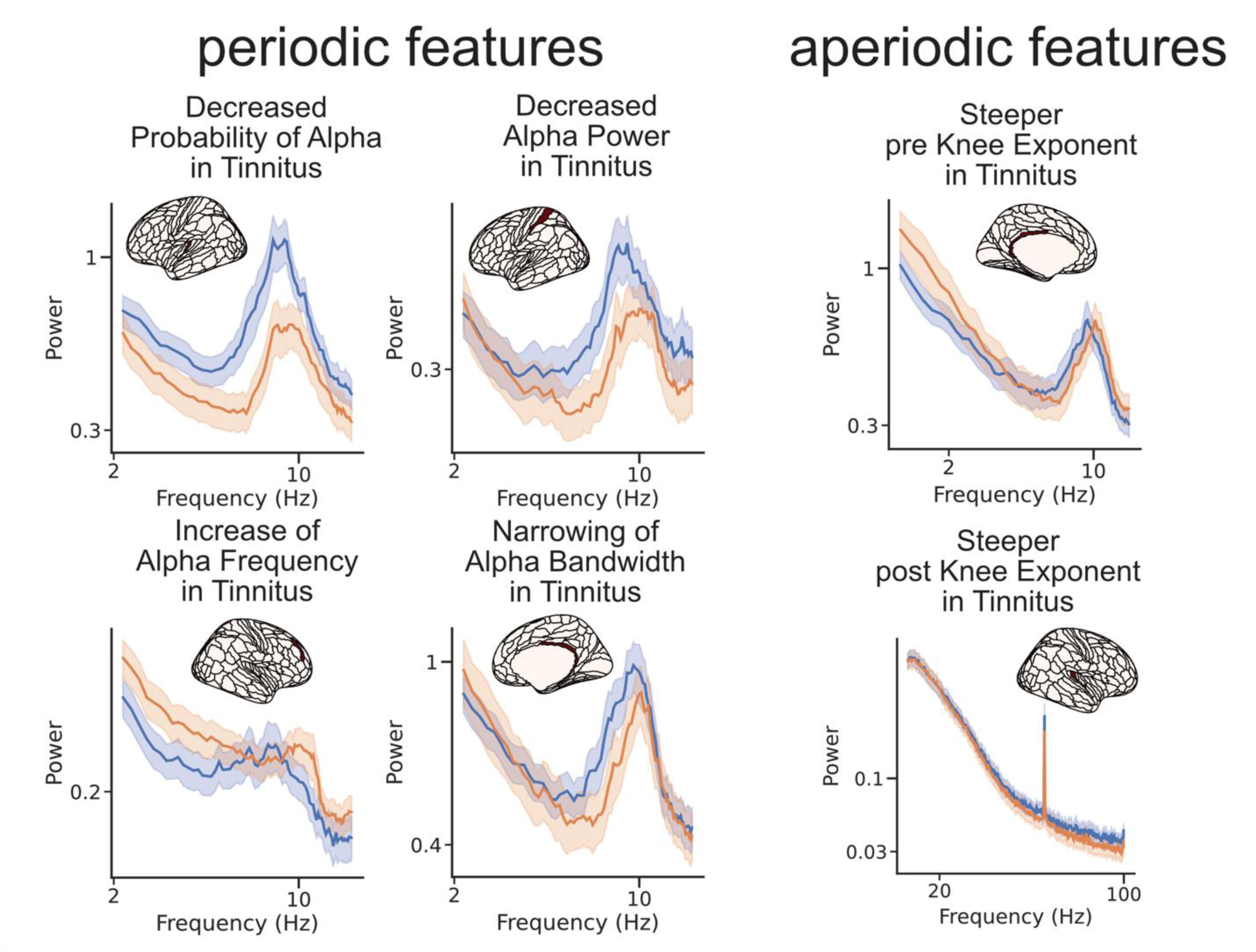
: **Selected raw power spectra from representative regions showing periodic and aperiodic effects.** Power spectra from regions within the corresponding clusters identified in Figure 1 are shown to illustrate the spectral patterns underlying several of the observed periodic and aperiodic effects. Shaded bands indicate the standard error of the mean calculated across subject within the parcel highlighted in red.

**Supplementary Figure S3.**
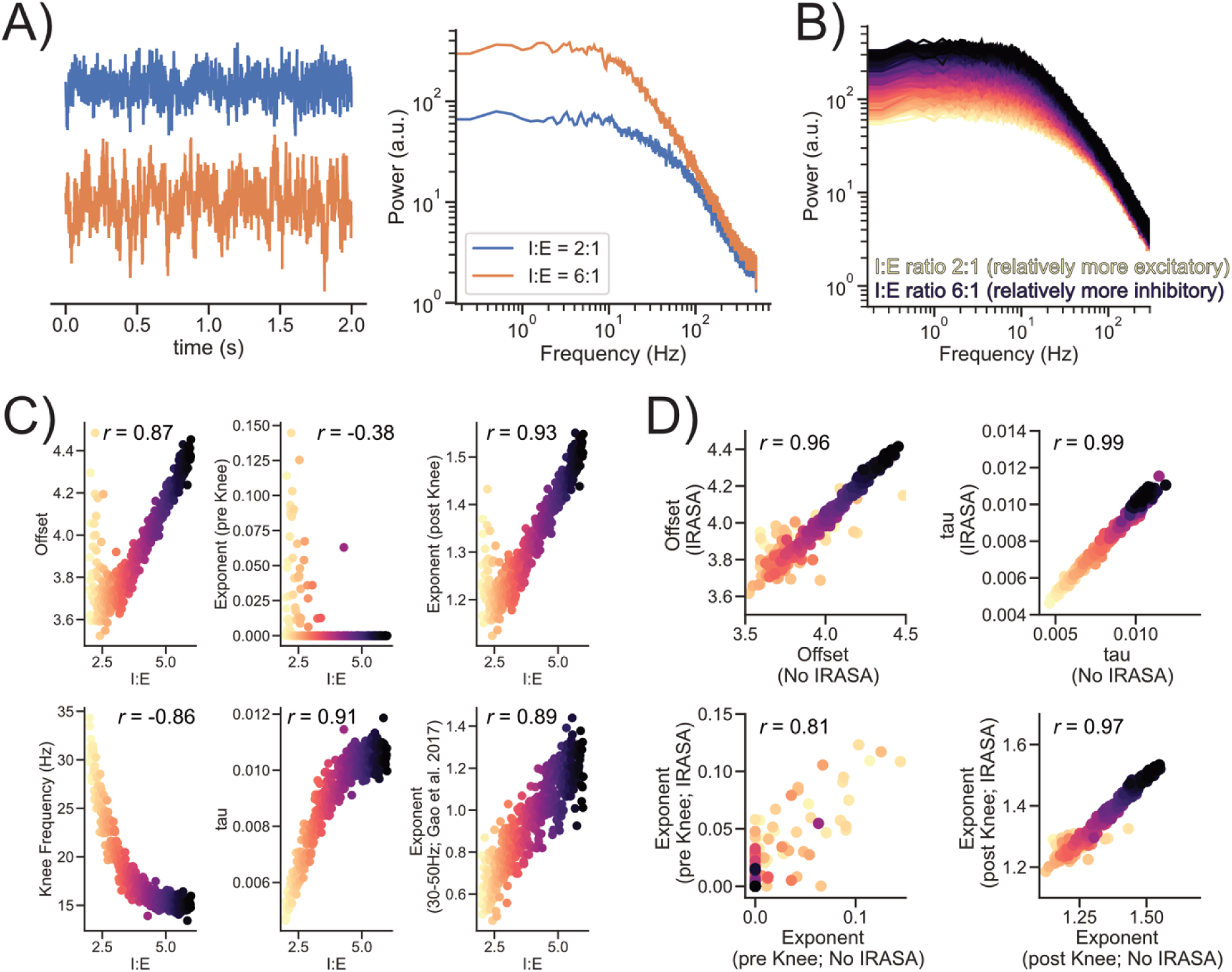
: Simulated neural time series based on the model of Gao et al. 2017 [37], illustrating effects of changes in inhibition:excitation ratio (I:E). **A)** Neural time series were simulated using the parameters reported by Gao et al. (2017) [37] for I:E ratios ranging from 2:1 to 6:1. Example time series and corresponding power spectra are shown for I:E = 2:1 and I:E = 6:1. **B)** Power spectra spanning the full range of simulated I:E ratios were computed and parameterized using IRASA, implemented in PyRASA (Schmidt et al., 2025). **C)** Aperiodic spectra were further parameterized using the model described in *Methods – Spectral Analysis*. Multiple extracted parameters tracked the simulated changes in I:E, with association strengths similar to those reported for the 30–50 Hz exponent fit in Gao et al. (2017) [37]. **D)** Because IRASA can yield biased estimates in the presence of a spectral knee [27], we compared parameter estimates derived with and without IRASA. Offset, tau, and post-knee exponent estimates were highly consistent across approaches, whereas pre-knee exponent estimates showed evidence of bias.

**Supplementary Figure S4:**
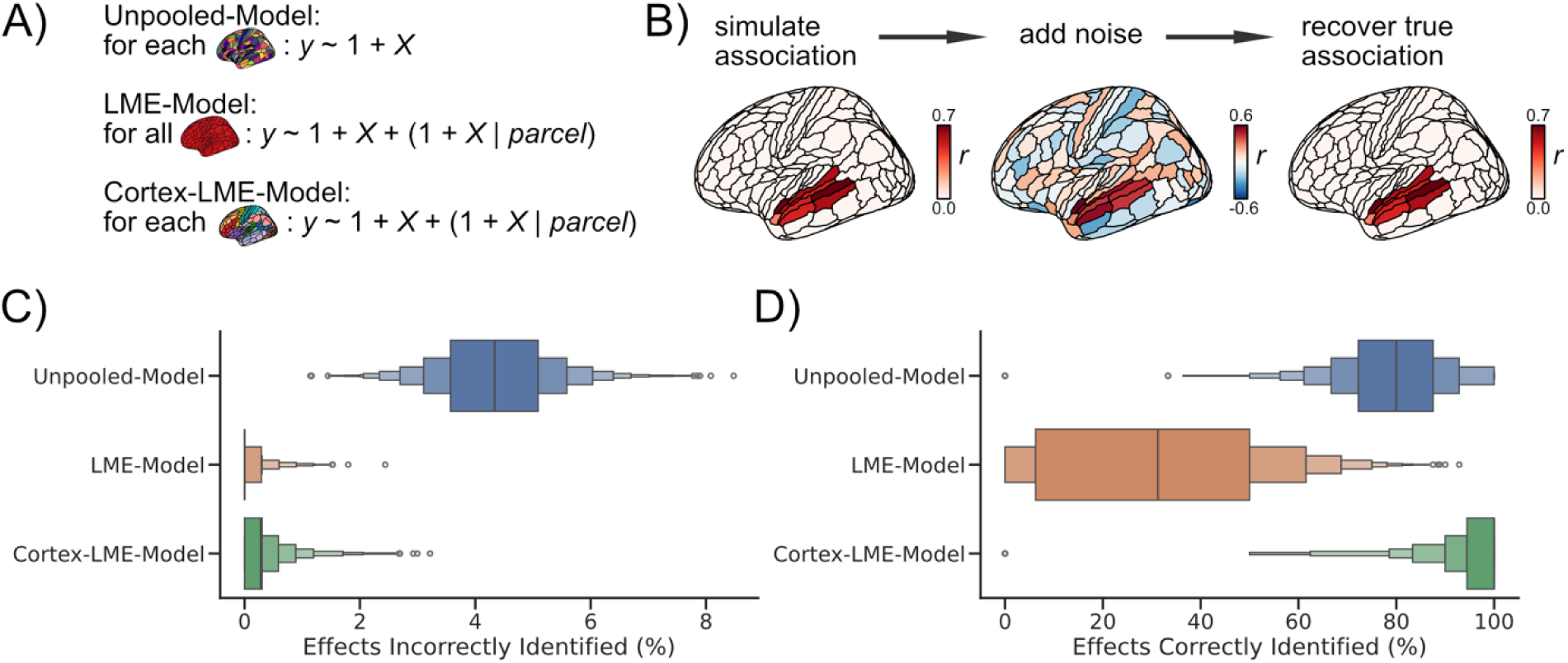
Sensitivity analysis of candidate Bayesian mixed-effects models. **A)** We evaluated three plausible model structures for relating a predictor variable (e.g., tinnitus) to neural activity: an Unpooled Model fit separately to each unit, a standard LME Model with partial pooling across parcels, and a Cortex-LME Model with partial pooling within cortical subdivisions. These models were compared on simulated data to determine which best matched our assumptions about MEG data. **B)** Because MEG effects are expected to be spatially clustered within cortical areas, we simulated standardized beta coefficients that varied between 0.3 and 0.7 across functionally similar neighboring parcels. Gaussian noise (*Normal*(loc=0, *SD*=1)) was added to these effects, and each model was then used to recover the true underlying associations. **CD)** The Cortex-LME Model achieved the best balance between sensitivity and false-positive control, with the highest proportion of correctly identified effects and the lowest proportion of incorrectly identified effects, justifying its use in the present study.

